# Efficient enrichment cloning of TAL effector genes from Xanthomonas

**DOI:** 10.1101/256057

**Authors:** Tuan Tu Tran, Hinda Doucouré, Mathilde Hutin, Boris Szurek, Sébastien Cunnac, Ralf Koebnik

**Affiliations:** IRD, Cirad, University of Montpellier, IPME, Montpellier, France

**Author notes:** Corresponding author: R. Koebnik.

**Keywords:** Enrichment cloning, Type III secretion, Type III effector, TAL effector, TALome, *Xanthomonas*, Bacterial leaf blight, Bacterial leaf streak, Rice

## Abstract

Many plant-pathogenic xanthomonads use a type III secretion system to translocate Transcription Activator-Like (TAL) effectors into eukaryotic host cells where they act as transcription factors. Target genes are induced upon binding of a TAL effector to double-stranded DNA in a sequence-specific manner. DNA binding is governed by a highly repetitive protein domain, which consists of an array of nearly identical repeats of ca. 102 base pairs. Many species and pathovars of *Xanthomonas*, including pathogens of rice, cereals, cassava, citrus and cotton, encode multiple TAL effectors in their genomes. Some of the TAL effectors have been shown to act as key pathogenicity factors, which induce the expression of susceptibility genes to the benefit of the pathogen. However, due to the repetitive character and the presence of multiple gene copies, high-throughput cloning of TAL effector genes remains a challenge. In order to isolate complete TAL effector gene repertoires, we developed an enrichment cloning strategy based on

- genome-informed *in silico* optimization of restriction digestions,
- selective restriction digestion of genomic DNA, and
- size fractionation of DNA fragments.

Our rapid, cheap and powerful method allows efficient cloning of TAL effector genes from xanthomonads, as demonstrated for two rice-pathogenic strains of *Xanthomonas oryzae* from Africa.

**METHOD NAME:** TAL fishing

## METHOD OVERVIEW

In previous studies, a few *tal* genes were isolated from African strains of *Xanthomonas oryzae*. Among them, only *tal5* from MAI1 and *talC* from BAI3 have been characterized as major virulence TAL effectors [1, 2]. These studies relied on screening of genomic DNA cosmid libraries to isolate and sequence *tal* genes, which is a laborious and time-consuming process. Due to the presence of multiple, very similar *tal* genes in *X. oryzae* strains and due to their highly repetitive character, PCR amplification is not feasible [3]. Recently, a method based on size fractionation of restriction-digested genomic DNA was developed, which made use of two conserved BamHI restriction sites, one at the ATG start codon and another one approximately 150 bp upstream of the stop codon [4]. Size-fractionated BamHI fragments (1.5 to 7.5 kb), covering most of the *tal* gene sequence, were cloned into pUC19, followed by transformation into *Escherichia coli*. Dot blot hybridization revealed that 115 out of 3000 clones contained a *tal* gene (i.e. less than 4%), which was further confirmed by Southern blot analyses and Sanger DNA sequencing. Here, we improve this method by making use of the extreme similarity among repeat DNA sequences and the strong conservation of the N- and C-terminal regions to identify frequently cutting restriction enzymes that do not cut within any of the *tal* genes. Complete combinatorial digestion of genomic DNA with BamHI and two additional frequent cutters for counter-selection, followed by size fractionation of DNA fragments allows rapid, cheap and efficient cloning of *tal* gene BamHI fragments from xanthomonads, as demonstrated with two African strains of *X. oryzae*.

### *In silico* combinatorial restriction digestion

Using BioEdit (<u>http://www.mbio.ncsu.edu/bioedit/page2.html</u>), we identified 54 restriction enzymes that would not cleave in any of 71 *tal* gene BamHI fragments from *X. oryzae* that were retrieved from GenBank. From this analysis, two restriction enzymes, ApaLI (GTGCAC) and SfoI (GGCGCC), were selected for further analyses. Genome sequences of nine *X. oryzae* strains were then used for *in silico* combinatorial restriction digestion using Microsoft Office software (Microsoft Word and Microsoft Excel) (Table 1). First, each genome sequence was converted into a consecutive list of BamHI fragments in Word. Upon conversion from text to table format, all virtual DNA fragments were transferred to an Excel spreadsheet. In Excel, fragment sizes and absence/presence of ApalI and SfoI sites were scored using appropriate formulas. DNA fragments were sorted by the absence/presence of the two restriction sites and the size of virtual BamHI fragments. *tal* gene-related BamHI fragments were identified by TBLASTN searches. Finally, data were re-analyzed assuming isolation of 2 to 5 kb DNA fragments prior to cloning.

**Table 1.** |In *silico* digestion analysis of *Xanthomonas oryzae* genome sequences.

| Clade | African *X. oryzae* pv. *oryzae* |  |  | Asian *X. oryzae* pv. *oryzae* |  |  |  | *X. oryzae* pv. *oryzicola* |  |
| --- | --- | --- | --- | --- | --- | --- | --- | --- | --- |
| Origin | Mali | Burkina Faso | Cameroon | Philippines | Philippines | Korea | Japan | Philippines | Burkina Faso |
| Strain | MAI1 | BAI3 | AXO1947 | PXO86 | PXO99<sup>A</sup> | KACC 10331 | MAFF 311018 | BLS256 | BAI11 |
| Accession number | pending | pending | CP013666 | CP007166 | CP000967 | AE013598 | AP008229 | CP003057 | CP007221 |
| Sequencing technology | PacBio | PacBio | PacBio | PacBio | Sanger | Sanger | Sanger | Sanger | PacBio |
| Reference | unpublished | unpublished | Huguet-Tapia *et al.*, 2016 | Booher *et al.*, 2015 | Salzberg *et al.*, 2008 | Lee *et al.*, 2005 | Ochiai *et al.*, 2005 | Bogdanove *et al.*, 2011 | Booher *et al.*, 2015 |
| BamHI | 769 | 756 | 750 | 827 | 883 | 815 | 829 | 845 | 885 |
| ApaLI | 1671 | 1651 | 1637 | 1596 | 1662 | 1589 | 1587 | 1672 | 1693 |
| SfoI | 4986 | 4959 | 4881 | 4954 | 5223 | 4941 | 4980 | 4165 | 4564 |
| Total<sup>1</sup> | 102 | 102 | 103 | 108 | 123 | 105 | 109 | 145 | 157 |
| 2-5 kb<sup>2</sup> | 15 | 15 | 15 | 23 | 28 | 21 | 23 | 31 | 27 |
| Number of *tal* gene BamHI fragments<sup>3</sup> | 9 | 9 | 9 | 16 | 18 | 13 | 16 | 27 | 23 |
| Number of *tal* genes<sup>4</sup> | 9 | 9 | 9 | 16 (2) | 18 (1) | 13 | 16 (1) | 27 (1) | 23 (1) |
| Percentage (BamHI fragments) | 1.2 | 1.2 | 1.2 | 1.9 | 2.0 | 1.6 | 1.9 | 3.2 | 2.6 |
| Percentage (Total) | 8.8 | 8.8 | 8.7 | 14.8 | 14.6 | 12.4 | 14.7 | 18.6 | 14.6 |
| Percentage (2-5 kb) | 60.0 | 60.0 | 60.0 | 69.6 | 64.3 | 61.9 | 69.6 | 87.1 | 85.2 |

<sup>1</sup> Number of BamHI-fragments with BamHI-compatible overhangs predicted after combinational digestion with BamHI, ApaI and SfoI
<sup>2</sup> Number of BamHI-fragments in the size range of 2 to 5 kb with BamHI-compatible overhangs predicted after combinational digestion with BamHI, ApaI and SfoI
<sup>3</sup> Number of *tal* gene-related BamHI fragments predicted to be cloned following combinatorial digestion
<sup>4</sup> Number of *tal* genes in the TALomes as published or predicted, including truncated *tal* genes (in brackets)

Assuming that only DNA fragments flanked by two BamHI overhangs would be cloned, but none of the fragments that contain only one BamHI overhang (which could nevertheless occur by head-to-tail ligation of two fragments flanked by one BamHI overhang and one overhang from the other two enzymes), one would expect to obtain between 10% and 20% of *tal* gene BamHI fragments without size fractionation and between 50% and 90% of *tal* gene BamHI fragments with size fractionation (2 to 5 kb DNA fragments). However, our simulation revealed that this procedure would miss to isolate *tal*-gene related DNA fragments from truncated *tal* genes (two in PXO86, one in KACC10331, one in BAI11), which nevertheless are of functional relevance [5, 6]. Most importantly, this method should allow isolation of BamHI fragments from all full-length *tal* genes, corresponding to the full functional TALome that induces the expression of resistance or susceptibility genes.

### Bacterial strains, plasmids and growth conditions

The bacterial strains used in this study were *Escherichia coli* DH5a (Stratagene, La Jolla, CA, USA) and *X. oryzae* strains BAI3 and MAI1 [7]. *E. coli* bacteria were cultivated at 37 °C in lysogenic broth (LB), *X. oryzae* strains at 28 °C on PSA medium (10 g peptone, 10 g sucrose, 1 g glutamic acid, 16 g agar, l^-1^ H_2_O). Antibiotics were used at the following concentrations: gentamicin, 20 µg/ml.

Plasmids were introduced into *E. coli* by electroporation and into *X. oryzae* by biparental conjugation using *E. coli* strain S17-1. The plasmid used to clone *tal* gene BamHI fragments, pSKX1, was obtained upon digestion of pSXK1-*talC* [2] with BamHI and re-ligation, leading to a construct where the translational start codon with an overlapping BamHI site (AT<u>GGATCC</u>, BamHI site underlined) of *talC* is fused to 151 base pairs corresponding to the 3’ end of the *talC* open reading frame downstream of the second BamHI site of *talC*.

### DNA extraction

Total genomic DNA of two African *X. oryzae* strains was prepared using the midi-prep Qiagen^®^ genomic DNA preparation protocol for bacteria using Genomic-tips 100/G (QIAGEN SAS, Courtaboeuf, France). The total genomic DNA was resuspended in 500 µl of double-destilled H_2_O, aliquoted into several Eppendorf tubes and stored at -30 °C. Plasmid DNA wase extracted using QIAprep Spin Miniprep Kit or QIAGEN Plasmid Kit for midiprep (Qiagen SAS).

### Enrichment cloning of *tal* gene BamHI fragments

20 μg of genomic DNA were digested with BamHI-HF, SfoI and ApaLI in CutSmart^®^ Buffer (New England Biolabs SAS, Enry, France) at 37 °C overnight. The High-Fidelity version of BamHI was used to avoid star activities upon prolonged incubation. Digestion products were purified using QIAquick PCR Purification Kit (Qiagen SAS) for column purification or using the QIAquick Gel Extraction Kit (Qiagen SAS) to isolate 2-5 kb DNA fragments. DNA fragments were cloned into BamHI-digested pSKX1, which was dephosphorylated by TSAP thermosensitive alkaline phosphatase (Promega, Charbonnières-les-Bains, France) and purified using the QIAquick PCR Purification Kit (Qiagen SAS). Upon ligation with T4 DNA ligase (Promega) at 4 **°**C overnight, DNA was transformed into *E. coli*. Transformants were plated onto LB-gentamycin plates and bacterial colonies were screened by PCR for the presence of a cloned *tal* gene BamHI fragment by polymerase chain reaction (PCR).

### PCR screening of bacteria for cloned *tal* gene BamHI fragments

To assess the presence and orientation of cloned *tal* gene BamHI fragments, two primers pairs were used in PCR assays. Firstly, primers pthXo1-nt-Fw1 (5’-GCAGCTTCAGCGATCTGCTC)and PthXo1-nt-rev2(5’-TCAGGGGGGCACCCGTCAGT) were used to amplify a DNA fragment of approximately 590 to 660 bp corresponding to the highly conserved N-terminal region of all *X. oryzae* TAL effectors (size variation is due to an in-frame deletion in a few *tal* genes). Amplification was carried out with an initial denaturation step of 5 min at 95 °C, 30 cycles of 30 s at 95 °C, 30 s at 62 °C, and 45 s at 72 °C, and a final elongation step of 10 min at 72 °C. Secondly, to determine the orientation of the cloned *tal* gene BamHI fragment in pSKX1, primer AvrXa7-Ct-Fw2 (5’-GCGTTGGCCGCGTTGACCAA), which anneals to the 3’ end of the cloned *tal* gene BamHI fragment, and pSKX1-Rev (5’-gggcaccaataactgcctta-3’), which anneals to the plasmid vector pSKX1, were used to amplify a DNA fragment of approximately 904 bp when a *tal* gene BamHI fragment is inserted in frame with the 3’ portion of the vector-borne *talC* fragment. Amplification was carried out with an initial denaturation step of 5 min at 95 °C, 30 cycles of 30 s at 95 °C, 30 s at 64 °C, and 1 min at 72 °C, and a final elongation step of 10 min at 72 °C. The empty vector, pSKX1, and a plasmid containing the *tal5* gene from strain MAI1, pSKX1-*tal5*, served as negative and positive control, respectively.

### DNA sequencing of cloned *tal* gene BamHI fragments

Plasmid DNA from positive colonies in the first PCR assay was digested with BamHI to estimate the insert size and thus, the expected number of repeats, via gel electrophoresis in a 1.2 % agarose gel. Repeat regions were Sanger sequenced from both sides with two oligonucleotide primers: forward (5’-GCCGGATCAGGGCGAGATAACT) and reverse (5’-CACTGACGGGTGCCCCCCTGAA). Plasmid DNA from in-frame clones, as revealed by the second PCR assay, were Sanger sequenced from both sides of vector using primers pSKX1-For (5’-GGCACGACAGGGTTTTCCCGAC) and pSKX1-Rev (5’-GGGCACCAATAACTGCCTTA).

### Restriction fragment length polymorphism analyses

To confirm that the two counter-selection restriction enzymes, ApaLI and SfoI, do not cut within *tal* gene BamHI fragments, we performed restriction fragment length polymorphism (RFLP) analyses. 3 μg of single (BamHI) or triple (BamHI, ApaLI and SfoI) digested genomic DNA of the African *X. oryzae* strains BAI3 and MAI1 were electrophoretically separated on 0.8% agarose gels in 0.5 × TBE (Tris-Boris-EDTA) buffer at 25 V for 84 h at 4 °C. Upon DNA fragments were transferred to an Amersham Hybond-N+ membrane by Southern blotting (GE Healthcare, Vélizy-Villacoublay, France). A 663-bp fragment of the *tal5* gene was PCR-amplified and digoxigenin-labeled with DIG-High Prime (Roche Diagnostics, Meylan, France) using primers pthXo1-nt-Fw1 and PthXo1-nt-rev2, as described above.

### Efficiency of *tal* gene BamHI fragment enrichment cloning

As a proof-of-concept, we applied our strategy to two African strains of *X. oryzae*, BAI3 and MAI1 [7]. 20 μg of genomic DNA of MAI1 was digested with BamHI only or in combination with ApaLI/SfoI, respectively. The digestion products were aliquoted and subjected to two independent purification treatments, i.e. column (total fragments) and gel purification (2 to 5 kb fragments). Both sets of DNA fragments were ligated into plasmid pSKX1. As shown in Table 2, *tal* gene-harboring colonies were rarely detected in both single BamHI digestion treatments (yielding 0.6% and 0.8% for column purification and gel purification, respectively). Upon combinatorial digestion, column purification resulted in 10 *tal* gene-positive clones among 324 tested colonies (3.1%). Strikingly, counter-selection digestion combined with size fractionation resulted in 28 out of 89 *tal* gene-positive colonies (31.5%), as revealed by PCR screening. Cloned DNA fragments of all 28 positive colonies were sequenced from both sides using primers that anneal to the plasmid vector, thus confirming the presence of *tal* gene BamHI fragments in all *tal* gene-positive colonies. Similar results were obtained when the enrichment cloning was repeated for strain MAI1 (25.0% *tal* gene-positive colonies) and applied to strain BAI3 (26.9% *tal* gene-positive colonies) (Table 2), thus demonstrating superiority of the new method over previous cloning approaches.

**Table 2.** |Efficiency of cloning of *tal* gene BamHI fragments from *X. oryzae* strains MAI1 and BAI3 upon different enrichment treatments.

| Treatment |  |  |  |  | Number of different cloned *tal* gene BamHI fragments |
| --- | --- | --- | --- | --- | --- |
| BamHI digestion | + | + | + | + |  |
| Gel purification of 2-5 kb DNA fragments |  | + |  | + |  |
| Double digestion with two counter-selection enzymes |  |  | + | + |  |
| **Strain MAI1** | 5 / 790<sup>a</sup> | 5 / 630 | 10 / 324 | 28 / 89 & 34 / 136 | 9 |
|  | 0.6%<sup>b</sup> | 0.8% | 3.1% | 31.5% & 25.0% |  |
| **Strain BAI3** | nt | nt | nt | 57 / 212 | 8 |
|  |  |  |  | 26.9% |  |

<sup>a</sup> Number of colonies with cloned *tal* gene BamHI fragments / number of analyzed colonies
<sup>b</sup> Percentage of colonies with cloned *tal* gene BamHI fragments
+ indicates that the sample was treated as indicated on the left.
nt, not tested

## Additional information

### Background information and significance of the method

Strains of *Xanthomonas* spp. cause important diseases of many economically important crop and ornamental plants. In most cases, pathogenicity depends of a set of type III effector proteins, which are injected into host cells via a molecular syringe, the type III secretion system (T3SS). Among all the type III effectors, one class is of particular interest for *Xanthomonas*: the Transcription Activator-Like (TAL) effectors. TAL effectors are conserved in many *Xanthomonas* spp and some of them have been shown to significantly contribute to pathogenicity [8]. Upon injection via the T3SS, TAL effectors localize into the host nucleus to directly or indirectly activate the expression of specific host genes [9, 10]. Among them are so-called susceptibility (*S*) genes, the induction of which promotes bacterial colonization in the affected plant tissues and/or development of disease symptoms [11].

Plant gene induction by TAL effectors depends on their central repeat region as well as C-terminally located nuclear localization signals and an activation domain [8]. The central repeat region is composed by almost identical tandem repeats (typically ranging from 33 to 35 amino acids) where residues at positions 12 and 13 are hypervariable, also referred to as repeat variable di-residues (RVDs). As demonstrated by the TAL effector-DNA binding code, the string of RVDs of each TAL effector determines the DNA sequence (or effector binding element, EBE) to which it binds [12, 13]. Upon refinement of the code, various *in silico* platforms were developed that allow prediction of TAL effector target genes in complex plant genomes [14, 15, 16]. In order to understand the collective function and evolution of TAL effector genes, there is a need to isolate and sequence complete repertoires of TAL effectors (e.g. TALomes) from multiple strains of *Xanthomonas*. Yet, efficient cloning was hampered by the facts that the genes have a highly repetitive structure and that most strains contain multiple copies of TAL effectors, thus limiting the usefulness of PCR-based approaches [3]. Recently, long-read, single molecule, real-time (SMRT), a.k.a. PacBio sequencing technology emerged as a new strategy for full TALome sequencing [17, 18, 19]. Yet, functional studies require the molecular cloning of *tal* genes. Here, we obtained the full TALome of strain MAI1 in only two weeks. Remarkably, this protocol can easily be parallelized and the obtained clones in the expression vector pSKX1 can directly be used in pathogenicity assays. Consequently, our new enrichment cloning procedure is expected to spur TALome research by allowing medium-throughput TALome cloning.

Despite the fact that the procedure was developed for strains of *X. oryzae*, it can easily be adapted to other xanthomonads as long as the sequence for a few *tal* genes is known, thus allowing to find appropriate restriction enzymes for counter-selection. Before use of new enzyme combinations it is recommended to perform Southern blot analyses comparing single and multiple digested DNA samples in order to avoid enzymes that would cleave within the *tal* gene BamHI fragments.

## Acknowledgements

We are grateful to thank Jonathan M. Jacobs for valuable comments.

